# Gene expression responses to environmental cues shed light on components of the migratory syndrome in butterflies

**DOI:** 10.1101/2024.07.17.602486

**Authors:** Daria Shipilina, Lars Höök, Karin Näsvall, Venkat Talla, Aleix Palahí, Elenia Parkes, Roger Vila, Gerard Talavera, Niclas Backström

## Abstract

Migration is a complex behavior involving the synchronisation of many physiological and behavioral processes. Environmental cues must thus be interpreted to make decisions regarding resource allocation between, for example, migration or reproduction. In butterflies, the lack of host plants to sustain a new generation may indicate the need to migrate. Here, we used the painted lady butterfly (*Vanessa cardui*) as a model to characterize gene expression variation in response to host plant availability. Assessment of the response to host plant availability in adult female butterflies revealed significant modifications in gene expression, particularly within hormonal pathways (ecdysone oxidase and juvenile hormone esterase). We therefore hypothesize that tuning the ecdysone pathway may play a crucial role in regulating the timing of migration and reproduction in adult female painted lady butterflies. In addition, our analysis revealed significant enrichment of genes associated with lipid, carbohydrate, and vitamin biosynthesis, as well as the immune response. As environmental acquisition occurs throughout the life cycle, we also tracked gene expression responses to two other environmental cues across major developmental stages. Differences in both larval crowding and host plant availability during development resulted in significant changes in the expression of genes involved in development, reproduction and metabolism, particularly at the instar V larval stage. In summary, our results offer novel insights into how environmental cues affect expression profiles in migratory insects and highlight candidate genes that may underpin the migratory syndrome in the painted lady butterfly.

## 1 INTRODUCTION

Animals are recurrently facing challenges to optimize resource allocation and individual decisions can have considerable downstream consequences on both survival and reproductive output (1,2). Migration is one example of a behavioral response to seasonal shifts in the environment, essentially allowing migratory organisms to avoid temporary unfavorable environmental conditions (3,4). Migratory movements have been characterized in detail in many different groups of organisms, however vertebrates have traditionally received most of the attention, while the understanding of invertebrate migration is limited to a few model species (5–7). Recent advances in tracking migratory movements in insects, for example via pollen metabarcoding and isotope-based geolocation of natal origins (e.g. 8–10), have revealed that they are capable of traversing remarkable distances. Migratory behavior is a complex trait involving decisions in initiating, maintaining and terminating migratory movements. Phenotypic plasticity in response to environmental cues is critical for making optimal decisions. Therefore, one of the crucial aspects for understanding the genetic basis of migration is pinpointing gene regulatory networks leading to the initiation of the migratory syndrome as a response to the environment (11). While environmental cues play a vital role in triggering behavioral switches, the underlying mechanisms of their interpretation and processing, as well as the responses on the molecular level, have only been studied in a few migratory insects e.g., the migratory locust *(Locusta migratoria)* and the monarch butterfly *(Danaus plexippus)* (12,13).

Upon sensing environmental cues, migratory insects commonly face scenarios demanding trade-offs between alternative resource allocations and physiological responses (14–16). The key trade-off characterizing the migratory syndrome, or the initial impulse to migrate, in insects is termed the oogenesis-flight syndrome and refers to the delayed investment in reproduction in favor of migration (17,18). Different migratory insect species exhibit significant variation in their physiological and behavioral integration of the oogenesis-flight syndrome. The syndrome varies from complete reproductive arrest during migration in for example the boll weevil (19) and the beet webworm (20), to the expression of the syndrome in certain generations like in the monarch butterfly (21), to an absence of reproductive arrest in several species, for example in the beet armyworm (22) and the codling moth (23), where females can migrate with fertilized and developed eggs (reviewed in 16). Most of the evidence for the oogenesis-flight syndrome is primarily based on phenotypic observations (18,24) and, while physiological and behavioral changes have been described in some detail, characterization of the genetic underpinnings of the trade-offs have predominantly been focused on reproductive arrest (25,26). Therefore, more detailed investigations are needed to enhance our understanding of this complex phenomenon.

Accurate perception of environmental cues is essential for the expression of the migratory syndrome, both in adult individuals and during ontogenesis (27). Among others, two environmental cues perceived during development have been shown to be associated with variation in propensity to migrate: rearing density and periodic starvation. High density during early developmental stages, for example, can lead to a predominant investment in migration, likely as a strategy to disperse from areas where competition with conspecifics is high. The desert locust *(Schistocerca gregaria)* is a notable example of this phenomenon, exhibiting a density-dependent phase polyphenism that triggers a transition from a benign, solitary phase to a more gregarious, highly migratory phase (12,28,29). In Lepidoptera, density-dependent migration has also been observed in the fall armyworm *Spodoptera frugiperda*; (30), and larval density has been associated with outbreaks in the agricultural pest, beet webworm *(Loxostege sticticalis* (20)). Food availability and quality has also been linked to the oogenesis-flight syndrome in insects, where limited resources during development predominantly manifest in reduced body size, fat storage, fecundity and investment in reproduction. Food availability should therefore have a major influence on migration capacity/propensity and, hence, the trade-off between reproduction and migration (31–33).

The painted lady butterfly *(Vanessa cardui)* is an emerging model species for studying the genomic basis of multi-generational long-distance migration (34–36). In addition to performing the longest individual migratory flight distances of any Lepidoptera (8,37–39), *V. cardui* is completely lacking diapause, which highlights the recurrent balance between reproduction and migration as a key adaptation in the species. Delayed onset of reproduction suggestive of an oogenesis-flight syndrome has been observed in *V. cardui* (40), but evidences are mixed (41) and considerable interindividual differences in migration distance have been observed (10). Recently developed genomic resources (35,36) make it possible to investigate the genetic underpinnings of migratory behavior in this species. Until now, however, very few attempts have been made to characterize the components of the migratory syndrome and its dependence on environmental cues. Spearheading work using methylation and chromatin accessibility data has pinpointed candidate pathways that are likely involved in sensory perception of environmental cues (42,43), but analyses that investigate potential associations with transcription profiles of specific genes or gene categories have not been performed.

Given the complex nature of genetic regulation underlying the migratory behavior, it is advantageous to conduct separate experiments targeting various environmental cues. Here, we make a first attempt to investigate the transcriptomic response to two environmental cues that can be associated with investment in reproduction or migration in butterflies: larval density and host plant availability for egg laying and as a food source. The main aims were to i) characterize transcriptomic responses to these environmental cues across developmental stages and in adult females, ii) identify developmental time points at which the environmental cues trigger a difference in gene expression, iii) identify candidate genes that might be involved in the trade-off between reproduction and migration.

## 2 MATERIALS AND METHODS

### 2.1 Experimental setup

Painted lady *(Vanessa cardui)* butterfly females were collected in Catalonia, Spain, and individually housed in cages for egg laying at 23°C under an 18:6-hour light:dark regime. The butterflies were provided with host plants *(Malva sylvestris)* for egg laying and a 10% sugar water solution as a food source. The F1 offspring were raised individually with *ad libitum* access to food plants *(M. sylvestris)*, and were subsequently divided into experimental groups under controlled environmental conditions (Figure 1).

**FIGURE 1.**
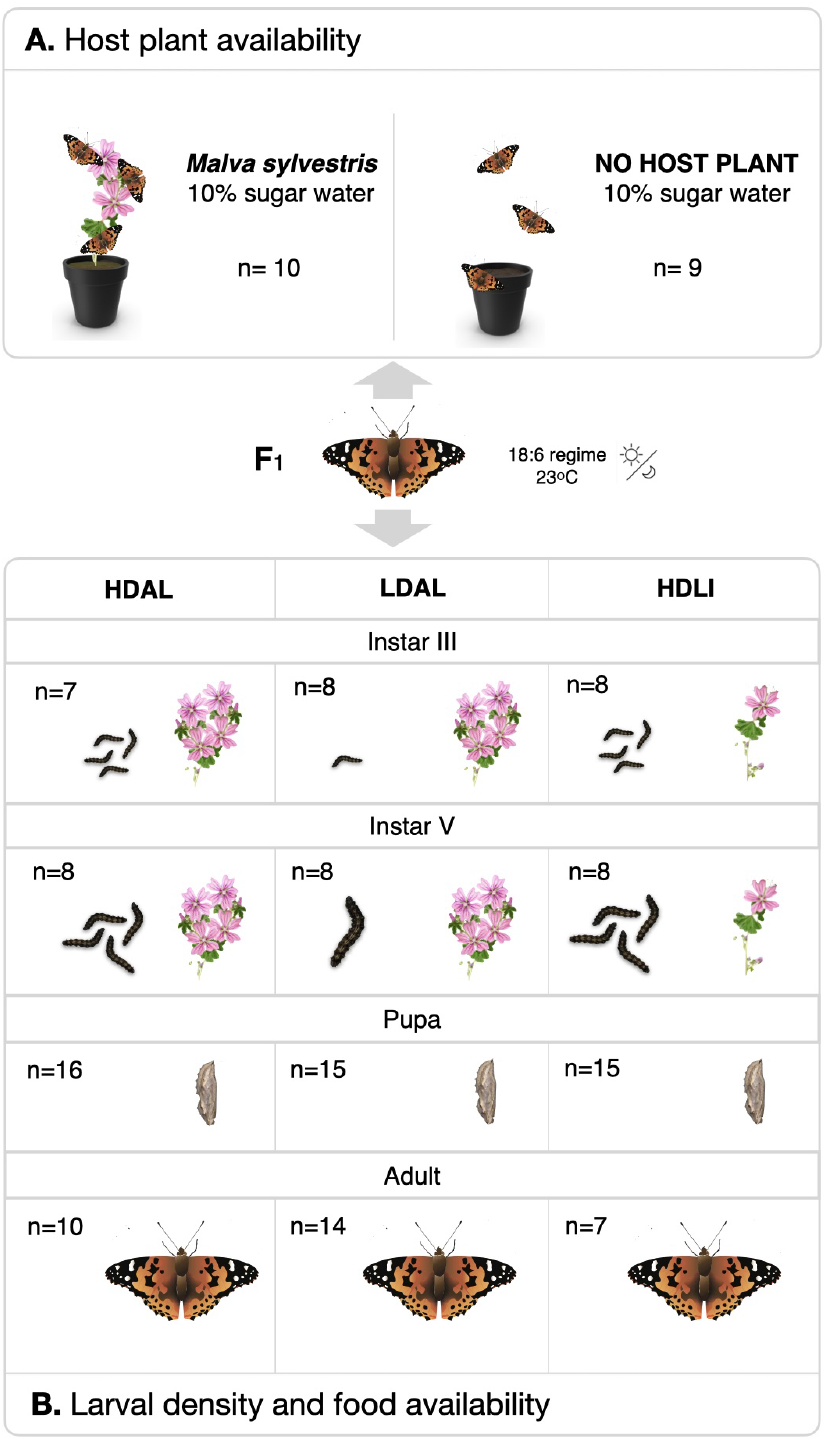
Setup of the two experiments conducted on offspring of wild-caught *Vanessa cardui* females (F1 in the center). Numbers of sequenced individuals are provided for each treatment and cohort. A) The host plant availability experiment, where recently emerged females were divided in two experimental groups with or without access to *Malva sylvestris* for egg laying. B) The setup of the larval density and food availability experiment for different developmental stages. Here, F2 offspring from five different F1 females were divided into three cohorts where the environmental conditions varied. HDAL = high density (10 larvae / flask) and *ad libitum* food, LDAL = low density (1 larva / flask) and *ad libitum* food, HDLI = low density (1 larva / flask) and limited food (fed every other day).

Two separate experiments were carried out to analyze the transcriptomic response to different environmental cues. The first experiment was designed to investigate the potential influence of the presence or absence of the host plants for egg laying on gene expression profiles in adult females (Figure 1A). The second experiment was designed to characterize the effects of larval density and food availability on gene expression profiles during development (larval instars III and V, and pupae) and after emergence (imagines, both sexes) (Figure 1B).

### 2.2 Host plant availability effects on gene expression profiles in adult females

Twenty newly emerged adult F1 females were marked individually and released into one of two large cages (80 × 80 × 50 cm), 10 in each cage. One cage contained an abundance of host plants (nine 15 × 15 cm pots with *M. sylvestris*) for egg laying, while the other cage lacked host plants (Figure 1A) and both cages contained 10 free-flying males. Both experimental groups were provided with 10% sugar water as adult food, and the temperature and light regime were the same as for rearing larvae (23°C and an 18:6-hour light-dark regime). In the morning five days after emergence, around the expected time for first mating / reproductive maturity (40,41), five females from each respective treatment group were snap frozen in liquid nitrogen.

### 2.3 Crowding and food availability effects on expression profiles across developmental stages

In the second experiment, five newly mated F1 females were placed in individual cages containing *M. sylvestris* for egg laying. The eggs (F2) laid by each female were collected and divided into three treatment groups (Figure 1): LDAL (low density, *ad libitum* food), HDAL (high density, *ad libitum* food), and HDLI (high density, limited food) (Figure 1B). In the LD (low-density) condition, larvae were individually reared in 1-liter flasks, while in the HD (high-density) treatment, 10 larvae were kept together in a single 1-liter flask. Both density treatment groups had *ad libitum* access to food *(M. sylvestris)*, which was replaced daily. In the LI (limited resource) treatment group, the food was replaced every other day, creating a mild starvation regime. Individuals in this treatment group were reared in groups of 10 (high density). This setup allowed us to contrast treatments with different food availabilities (HDAL versus HDLI) and larval rearing densities (HDAL versus LDAL) separately (Figure 1).

Samples were collected at four developmental stages: larva (instar III and instar V), pupa, and adult. Larvae were harvested on the day they entered the respective larval stage, pupae were sexed and collected one day after pupation, and adults were harvested in the morning on the day of emergence. Prior to RNA extraction, individuals were snap frozen in liquid nitrogen and stored at -80°C. Note that larvae of this species cannot be easily sexed, and these cohorts therefore can constitute a mix of males and females. Pupae and adults were sexed and divided into sex-specific cohorts. For each treatment and cohort, one individual among the offspring of each of the five different F1 females was selected for sequencing (Figure 1).

### 2.4 RNA extraction and sequencing

Two types of tissues were used for RNA extractions; heads (including antennae) and abdomens (the 6th - 8th body segments). The number of samples for each treatment/cohort are provided in Figure 1 and Supplementary Table 1). Tissues were homogenized using a micro-pestle in guanidine-isothiocyanate lysis buffer, followed by mixing with QiaShredder (Qiagen). RNA extractions were performed using the RNeasy Mini Kit (Qiagen) following the recommended guidelines by the manufacturer. RNA integrity and fragment lengths were assessed using 1% agarose gel electrophoresis, followed by measurements of the concentration using NanoDrop (ThermoFisher) and Qubit (ThermoFisher). Sequencing libraries were prepared using the Illumina TruSeq Stranded mRNA polyA selection kit and sequenced by the National Genomics Infrastructure (NGI) in Stockholm. Sequencing was conducted on two lanes of one S4 flow cell on the NovaSeq S6000 platform, generating 150 bp paired-end reads.

### 2.5 Differential expression analysis

For all steps of the read processing, from adapter filtering to read mapping and transcript quantification, the Nextflow nf-core (44) pipeline rnaseq v3.8.1 was applied (45). In brief, raw sequencing reads were trimmed using Cutadapt v.3.4 (46) as implemented Trim Galore! v0.6.7. STAR v2.7.10a (47) was used for mapping the reads to a previously published genome assembly (36). Read quantification was carried out using salmon v1.5.2 (48), and gene expression levels were measured in transcripts per million reads (TPM) values. Differential expression analyses were conducted in R v4.2.1 using DESeq2 v1.28.0 (49).

To assess differential expression between the cohorts of adult individuals with or without access to host plants for egg laying, we employed the Wald test in the DESeq2. Our experimental design incorporated the correction for potential family effects, with treatment as the primary variable (∼family+treatment). Due to incomplete family assignment for some samples, we utilized PCA analysis to recover the missing assignments. Additionally, we applied the Wald test for differential expression analysis in adult individuals subjected to environmental stressors: food limitation and larval crowding. Here, we also accounted for the potential effect of sex since both males and females were used in the analysis (∼family+sex+treatment).

Differential gene expression across developmental stages was assessed using the likelihood ratio test mode of DESeq2 (model = “LRT”), which allows for analysis of time course experiments. This test compared the fit of a full model (∼family+devstage+treatment+treatment:devstage) with a reduced model that excluded the interactive effect between the treatment and developmental stage (“devstage”) variables. This analysis aimed to evaluate whether the effect of the treatment on gene expression differed across developmental stages. The same model was applied to both head and abdomen tissues, and the analysis included the treatments food availability (HDAL versus HDLI) and rearing density (HDAL versus LDAL).

For further analysis, candidate genes were selected based on the criteria of an adjusted *p*-value < 0.05 (Benjamini and Hochberg to control FDR) and a log_2_ (fold change) > 2. The GeneOverlap package (50) was used to assess the significance of overlaps between candidate gene sets in different tissues. Since tests using LRT typically result in larger gene sets, clusterProfiler v3.17 (51) was applied to identify functional clusters within all sets of candidate genes across the four ontogenetic stages, using parameters (consensusCluster = TRUE, groupDifference = 2) on *r* log-transformed data. Regularized log (*r* log) transformation was used for stabilizing variance and normalizing the count data. In the case of the head tissue, where 745 candidate genes were identified, more stringent clustering parameters were employed with a group difference of 3. Functional information for differentially expressed genes was collected both by using previous annotations (35,36) and by BLAST searches against the entire nucleotide database (52). Data was summarized in corresponding supplementary tables, when two annotations were available the one from the most similar ortholog was chosen.

To assess if certain functional categories were overrepresented in the gene sets with significant differential expression, we conducted two types of enrichment analyses, Gene Ontology (GO) terms and KEGG pathways, utilizing previously obtained functional annotations. Enrichment analysis of GO terms (biological processes category) was performed using the TopGO package (53), employing the “weight01” algorithm, with a significance threshold of p < 0.01. The enrichment analysis of KEGG terms was performed using the enricher module of clusterProfiler (51).

## 3 RESULTS

### 3.1 Gene expression patterns in response to host plant availability in adult females

To understand how the presence or absence of host plants for egg laying affects transcriptional responses in recently emerged female imagines, we analyzed gene expression in head and abdominal tissues (Figure 2). The rationale behind choosing those particular tissues was that signaling cascades should be initiated in the head based on sensory perception of the environmental cues and that this may manifest in temporal differences in investment in reproduction and migration, which might be picked up by gene expression differences in the abdomen (where the gonads are located). In total, abdominal tissue was analysed in 10 individuals (five for each treatment) and head tissue in 9 individuals (five and four individuals from the treatments with and without access to host plants for egg laying, respectively). In the head, we found that 88 genes were significantly differentially expressed (p < 0.05 after FDR adjustment) between treatment groups. Of those, 34 genes (0.3% of all genes analysed) had higher and 44 had lower expression (0.4%) in the treatment with access to host plants compared to the treatment without host plants (Figure 2A). In the abdomen, the corresponding numbers were 70 differentially expressed genes; 44 (0.4%) with higher and 26 (0.2%) with lower expression in the treatment with access to host plants (Figure 2B). A list of the top significantly differentially expressed genes (p < 0.01) and their putative functions is provided in Supplementary Table 2.

**FIGURE 2.**
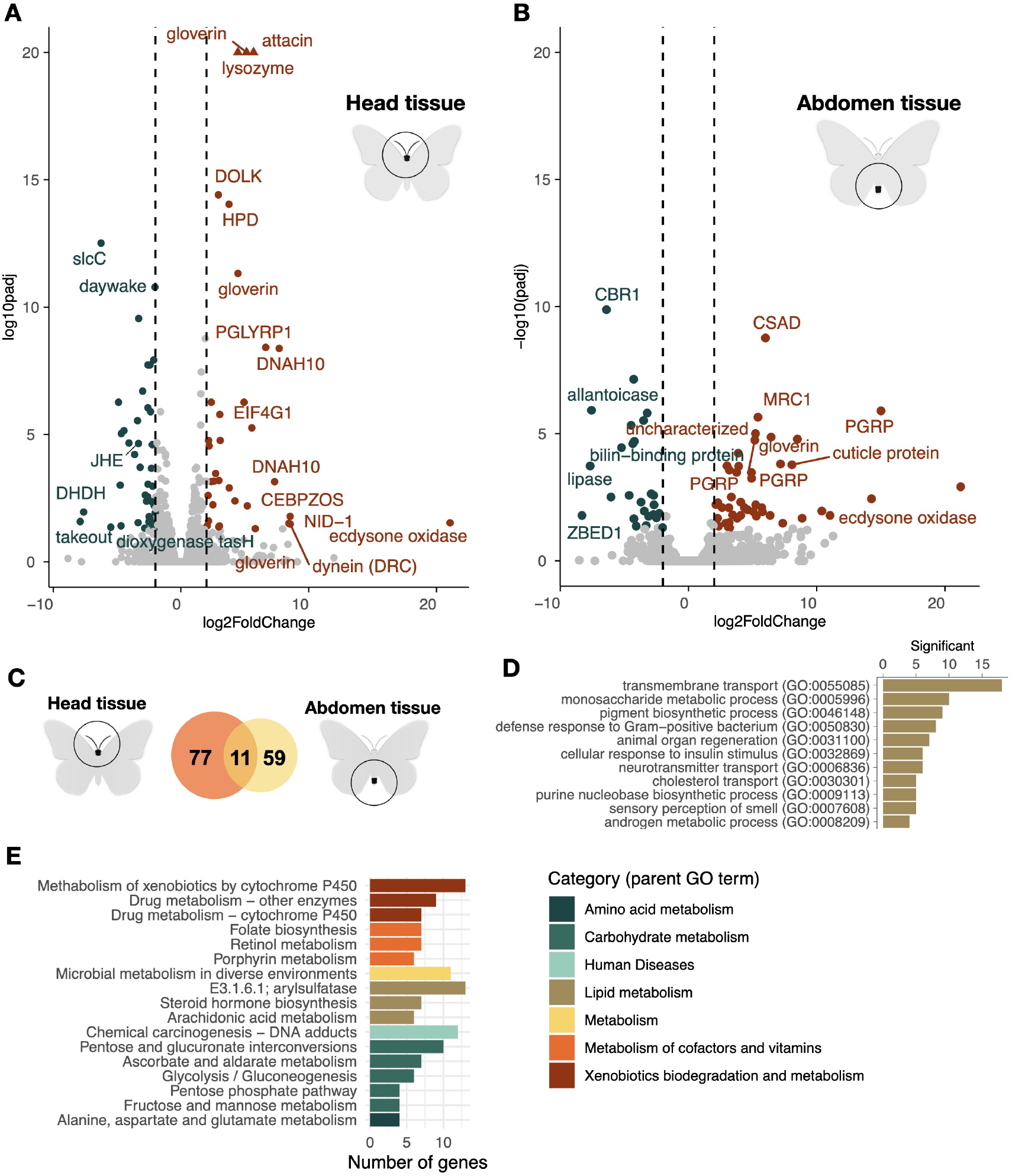
Volcano plots illustrating the relative levels of gene expression in the head (A) and abdomen (B) of adult females (x-axis; log_2_ fold change) in the two treatment groups with and without access to host plants for egg laying. Genes with a fold change difference > |2| and FDR-adjusted *p*-value < 0.05 are depicted in dark orange (significantly higher expression in the treatment with host plants) and teal (significantly higher expression in the treatment without host plants), respectively, while non-significant genes are illustrated by grey dots. Only selected outlier gene names are shown. Outlier genes with log_2_ fold change values exceeding y-limit are shown by triangles, exact values provided in the supplement. (C) Venn diagram showing the number of overlapping and unique differentially expressed genes between the two tissues (head = orange, abdomen = yellow). (D) Bar plot showing the counts of the enriched (FDR-adjusted *p* < 0.01) gene ontologies for the significantly differentially expressed genes between the treatment groups for both tissues combined. (E) Bar plot illustrating the numbers of significantly differentially expressed genes enriched for KEGG pathways, both tissues combined. The higher hierarchical grouping is indicated by bar colors (legend to the right).

We found that significantly differentially expressed genes encompassed a diverse range of functional categories (Figure 2A, B), including immune genes (gloverin, attacin, cytochrome p450, peptidoglycan recognition proteins), metabolic genes (lipase), and genes involved in endoskeleton formation (cuticle protein) (Figure 2, Supplementary Table 2). Of particular interest was the ecdysone oxidase gene, which exhibited a remarkable change in expression in both head and abdomen in the individuals that had access to host plants for egg laying (Figure 2A, B; Note that DESeq2 may sometimes exaggerate outliers (54)).

Among the genes that were differentially expressed between the adult female treatment groups, 11 genes were found in both tissues (Figure 2C). To gain further insights into the associations between functions of differentially expressed genes and increase statistical power, we combined the results from both tissues for GO term enrichment test. Significantly overrepresented GO terms encompassed transmembrane transport, various metabolic processes (including ecdysone biosynthesis), and defense response (Figure 2D, Supplementary Table 3). Consistent with this finding, the analysis of overrepresented KEGG pathways revealed enrichment of different metabolic pathways, in particular lipid, carbohydrate, vitamin, and xenobiotic metabolism (Figure 2E, Supplementary Table 4).

### 3.2 Gene expression variation associated with food availability during development

To complement the analysis in adult females, we focused on investigating differential gene expression across developmental stages in experimental cohorts exposed to environments that varied in food availability and rearing density. Again, we focused on the head and abdomen for the same reasons as indicated above. For the contrast between experimental groups with differences in food availability during development (HDAL versus HDLI), the likelihood ratio tests revealed 745 and 321 significantly differentially expressed genes (FDR-adjusted p-value < 0.05) in the head and abdomen, respectively. Notably, the two sets of genes with differential expression in the two respective tissues demonstrated a high degree of overlap (Jaccard index = 0.1, *p*-value = 9.6 × 10^−30^; Figure 3A). To check which stages contributed the most to the overall differences in expression patterns between treatment groups, we performed a clustering analysis which groups genes based on the expression patterns across developmental stages, facilitating the identification of genes with similar profiles and potential functional relationships. In head tissue, the most prominent cluster comprised 123 genes (16.5% of the differentially expressed genes in this tissue; Figure 3B). The majority of expression differences within this cluster were observed in instar III larvae. Similarly, in the abdominal tissue, 149 genes (46.4% of the differentially expressed genes) formed a distinct cluster. Genes within this cluster predominantly showed differential expression in instar V larvae (Figure 3C).

**FIGURE 3.**
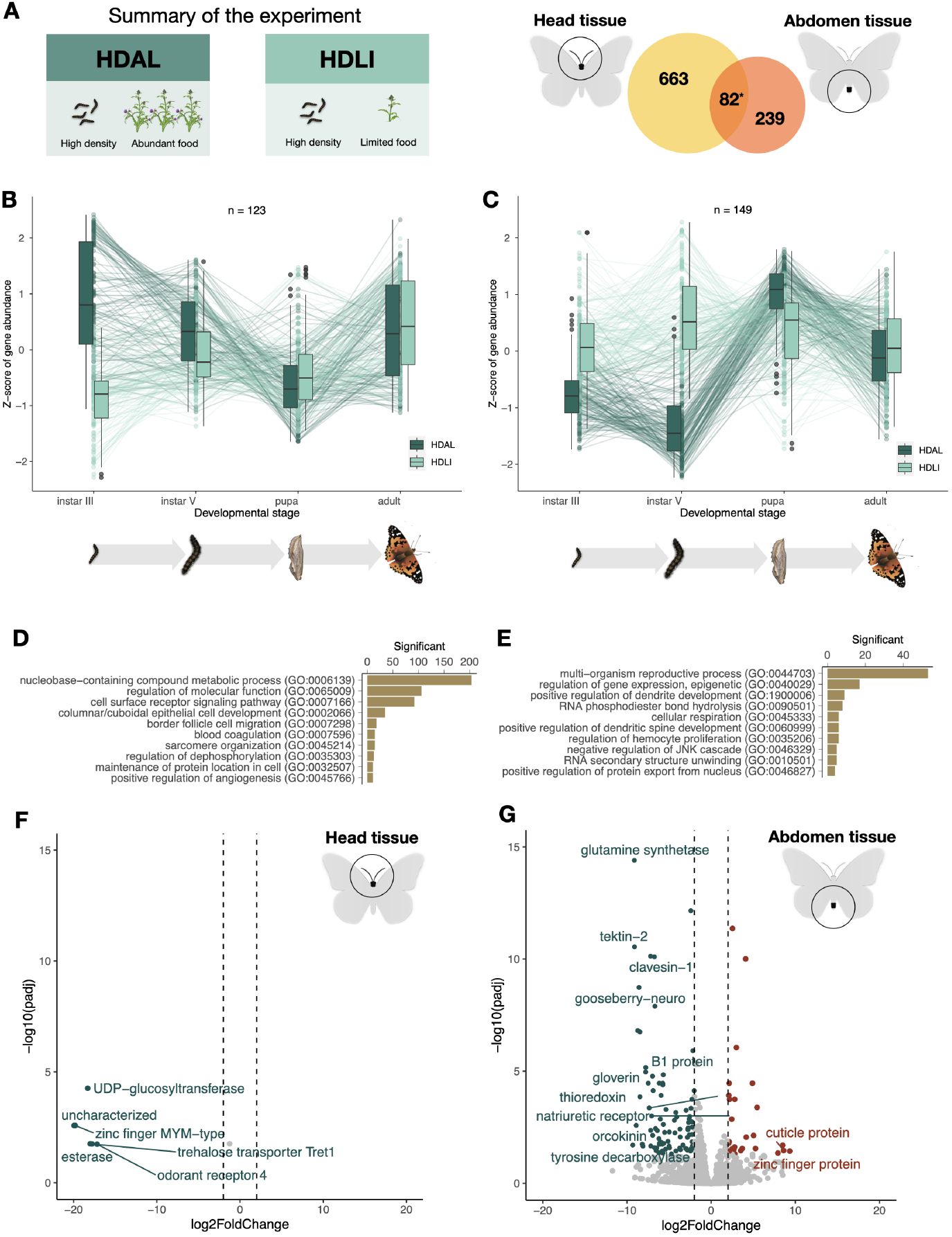
A summary of the results from the comparison between experimental groups with access to different amounts of food plants during development. (A) Summary of the experiment and a Venn diagram showing the number of differentially expressed genes in each respective tissue (left = head, right = abdomen) and the number of genes differentially expressed in both tissues (center). The star indicates that the number of overlapping genes was significantly higher than expected by chance. (B) and (C) Box plots showing the temporal patterns of differential expression across ontogenetic stages in the head (B) and abdomen (C). Outliers are indicated with circles, and temporal trends of gene expression levels for specific genes are illustrated with lines. (D) and (E) The top 10 most significantly overrepresented GO terms (p < 0.01) for differentially expressed genes in the head (D) and abdomen (E). (F) and (G) Volcano plots showing the relative levels of gene expression in the adult individuals for the head (F) and abdomen (G). Genes that are significantly differentially expressed and meet the threshold (FDR-adjusted *p*-value < 0.05, log_2_ fold change difference < |2|) are highlighted. Dark orange marks genes that are upregulated in adults in the food limitation treatment [HDLI], while teal spots mark genes that are upregulated in the treatment where larvae had access to unlimited food [HDAL].

The GO term analysis for differentially expressed genes in head tissue revealed both a general enrichment of functions related to metabolic processes and regulation and more specifically enrichment of functions associated with epithelial cell development, sarcomere organization, angiogenesis regulation and blood coagulation (Figure 3D, Supplementary Table 5). Differentially expressed genes in abdominal tissue were predominantly associated with reproductive processes, and neural and immune cell development (Figure 3E, Supplementary Table 6). In addition, a joint KEGG pathway analysis of both tissues unveiled that functions associated with ribosome biogenesis (ko03008) and aflatoxin biosynthesis (ko00254) were overrepresented.

The expression trajectories across developmental stages show that the influence of the environmental factors on gene expression differences between experimental groups, in general, appears to diminish at the pupal and adult stages. To investigate how environmental cues experienced during development are manifested in recently emerged imagines in more detail, we compared differences in gene expression between adult individuals that had experienced different environmental conditions during development specifically (Figure 3 F, G). In head tissue, only six genes (p < 0.05) showed significantly differential expression; *Tret1*, odorant receptor, UDP-glucosyltransferase, esterase and zinc-finger MYM (Supplementary Table 7). These genes were downregulated in response to limited food treatment. In the abdominal tissue, 189 (p < 0.05) genes were found to be differentially expressed between treatment groups. Among the most prominent outliers were cuticle protein (upregulated in response to limited food source), gloverin, glutamine synthetase, tektin, clavesin, gooseberry-neuro, orcokinin and tyrosine (downregulated in response to limited food source) (outliers are listed in Supplementary Table 7).

### 3.3 Gene expression variation associated with rearing density during development

To complement the analysis of gene expression variation associated with food plant availability during development, we also compared treatment groups that were reared at different densities (10 larvae versus 1 larva per flask, HDAL versus LDAL). In this comparison, we found a large number of genes differentially expressed in both the head (222 genes) and abdomen (372). There was also significant overlap between the tissues, i.e. a higher proportion of genes were differentially expressed in both tissues than expected by chance (Jaccard Index = 0.2, p-value = 1.2 × 10^−80^; Figure 4A).

**FIGURE 4.**
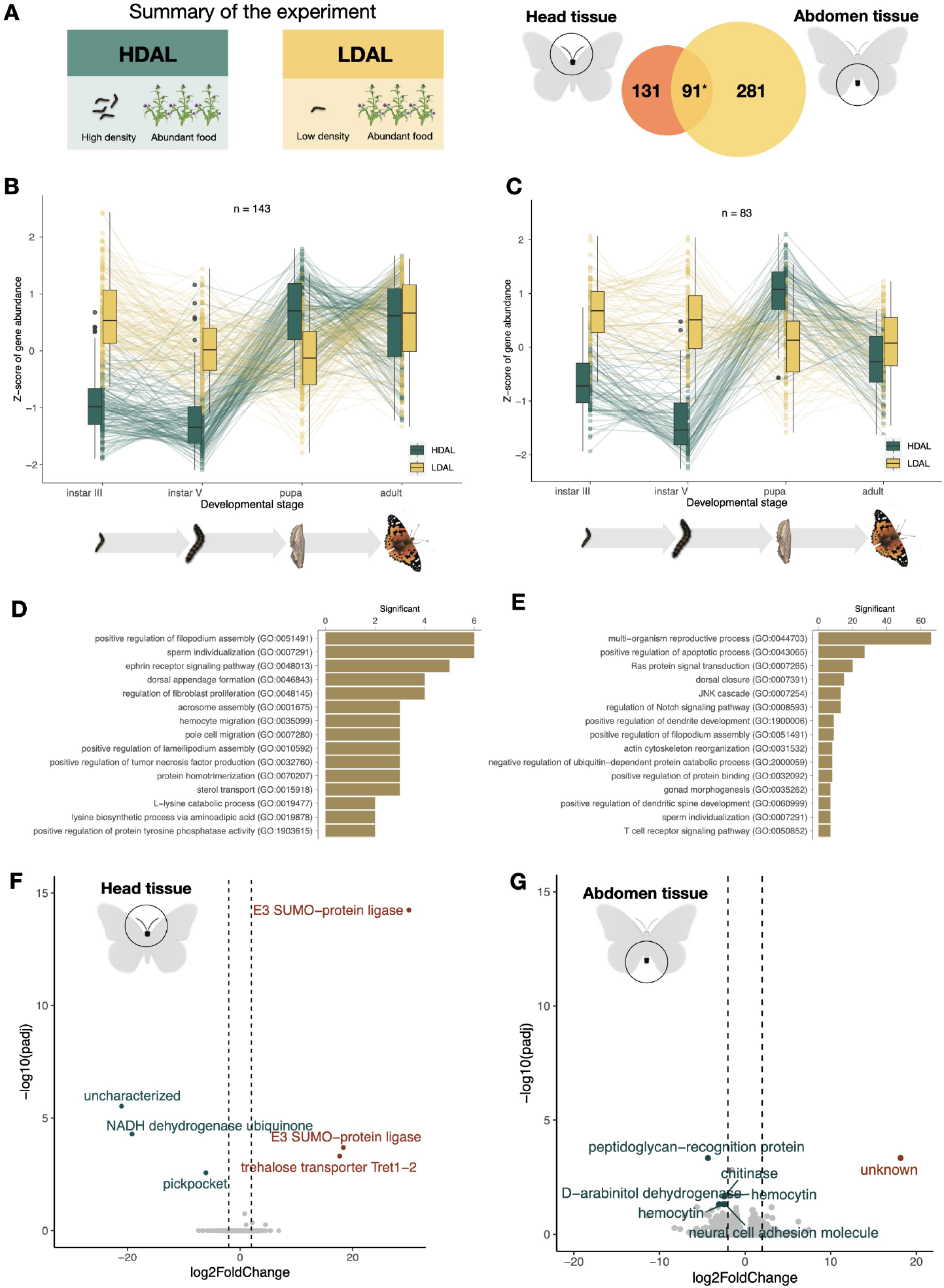
A summary of the results from the comparison between experimental groups, which were reared at different densities during development. (A) Summary of the experiment and a Venn diagram showing the number of differentially expressed genes in each respective tissue (left = head, right = abdomen) and the number of genes with significantly different expression in both tissues (center). The star indicates that the number of overlapping genes was significantly higher than expected by chance. (B) and (C) Box plots showing the temporal patterns of differential expression across ontogenetic stages in the head (B) and abdomen (C). Outliers are indicated with circles, and temporal trends of gene expression levels for specific genes are illustrated with lines. (D) and (E) The top 10 most significantly overrepresented GO terms (p < 0.01) for differentially expressed genes in the head (D) and abdomen (E), respectively. (F) and (G) Volcano plots showing the relative levels of gene expression in the adult female individuals for the head (F) and abdomen (G). Genes that are significantly differentially expressed and meet the threshold (FDR-adjusted *p*-value < 0.05, log_2_ fold change difference < |2|) are highlighted. Dark orange marks genes that are upregulated in adults in low density treatment [LDAL], while teal spots mark genes that are upregulated in the treatment where larvae had access to unlimited food [HDAL].

To investigate the temporal trajectories of differences in expression patterns between treatment groups during development, we again performed a clustering analysis based on the expression patterns across the different developmental stages. In head tissue, the two main clusters contained 143 (38.4% of the differentially expressed genes in this tissue; Figure 4B) and 69 (18.5%; Supplementary Feigure 1) genes, respectively. Visual inspection clearly showed that expression differences in instar III and V larvae were driving the overall patterns within these clusters (Figure 4B). In the abdominal tissue, 83 genes (37.4% of the differentially expressed genes in this tissue) formed a distinct cluster (Figure 4C). Again, this cluster was mainly distinguished by considerable differences in gene expression in the larval stages (Figure 4C).

The gene ontology enrichment analysis of functional roles of differentially expressed genes in these clusters revealed that the most enriched functional category was regulation of filopodia assembly in the head (Figure 4D). We also found an enrichment of ontology terms related to sperm maturation, ephrin signaling, and several other functional categories (Figure 4D, Supplementary Table 8). In the abdomen, there was a significant enrichment of GO terms associated with reproductive processes, including functions such as egg formation, egg laying, and mating development (Figure 4E, Supplementary Table 9). In addition, there were several enriched terms associated with signal transduction like ephrin signaling, Ras signal transduction (involved in cell growth, division, and differentiation), Notch signaling (associated with neurogenesis) and JNK signaling (regulation of ubiquitin-dependent processes).

Analogous to the analysis based on food plant availability, we compared differences in gene expression between recently emerged females that had experienced different levels of crowding during development (Figure 4 F, G, Supplementary Table 10). In this comparison, six genes showed significant differences in expression level in head tissue, of which functional information was available for five (Figure 4F). We found that two copies of the SUMO ligase and a trehalose transporter were significantly higher expressed in the low density (LDAL) compared to the high density treatment group (HDAL). The genes pickpocket and NADH dehydrogenase, in contrast, had a significantly higher expression in the HDAL than in the LDAL treatment group. In the abdominal tissue, the peptidoglycan recognition protein (*PGRP*), chitinase, hemocytin, D-arabinitol, and *NCAM* were significantly overexpressed in the LDAL treatment group compared to HDAL, while no genes with known functions had higher expression in HDAL.

## 4 DISCUSSION

### 4.1 Transcriptomic response to availability of host plants in adult female butterflies

The ability of individuals to switch between migration and reproduction is a key adaptation for migratory insects in general, and for long-distance migrants in particular (17–19). This capability can also be considered part of the migratory syndrome, which includes a suite of traits that facilitate migration (55). This study addresses the need to investigate the molecular mechanisms underlying the migration-reproduction trade-off, focusing on one of the environmental triggers for initiating or terminating migratory behavior in butterflies—the presence and abundance of host plants for egg-laying (40,56). We identified three major functional categories of genes activated in response to this treatment: hormonal regulation, metabolic regulation, and immunity. Below, we discuss these categories in detail and their connections to reproduction and migration.

Hormonal regulation has been shown to be important in controlling reproductive and migratory physiology and having a central role in the trade-off between reproduction and migration in butterflies (57). Similarly, our results from the analysis of gene expression variation associated with host plant availability for egg laying in adult females *V. cardui* underscore the importance of hormonal regulation for the plastic response to host plant abundance – i.e. we found significant changes in the expression of multiple genes regulating developmental hormones. Here, ecdysone oxidase stood out with a striking difference in expression level between experimental groups. Ecdysone is a steroid hormone crucial for numerous biological processes in metamorphic insects during major developmental transitions, including the maturation of oocytes and control of oviposition (58–60). Ecdysone oxidase in turn regulates the levels of active ecdysone by converting it to 3-dehydroecdysone and vice versa, thereby controlling the availability of active hormones through a rapid feedback mechanism. We propose that increased expression of ecdysone oxidase, triggered by the availability of host plants, modulates these hormone levels, enhancing reproductive investment (61).

Beyond ecdysone, other hormonal regulation pathways like juvenile hormone (JH) significantly contribute to the reproductive-migratory trade-off, as seen in monarch butterflies (25,62). Additionally, genes involved in JH synthesis were notably overexpressed in migrating hoverflies compared to sedentary ones (68). Similar patterns were observed in our study with genes such as juvenile hormone esterase (*JHE*; regulation of juvenile hormone levels; (62,64), and daywake and takeout genes (encoding juvenile hormone binding proteins), *nrf-6* (neuropeptide and hormone receptor), and cytochrome-P450 (control of ecdysone biosynthesis; (65)), suggesting a complex interplay of gene regulation and hormonal crosstalk crucial for adapting to environmental cues. We have also shown previously, using chromatin accessibility profiling, that *JHE* likely is upregulated in adult female butterflies with access to host plants for egg laying (43). Taken together, our observations corroborate that the ecdysone pathway and the regulation of juvenile hormone play pivotal roles in the plastic responses to environmental cues in insects in general (66,67) and that they may constitute key components in the trade-off between migration and reproduction in butterflies in particular.

The role of immunity in migratory syndrome is multifaceted, suggesting energy may be redirected from immune functions to aid migration or enhanced in response to varied pathogens (68,69). Our study revealed a strong immune response in adult females exposed to environments without host plants, evidenced by a large number of upregulated genes. Functional data from a diverse range of candidate genes identified in our study, allows us to speculate on the exact pathways of this response. Notably, multiple peptidoglycan-recognition proteins (*PGRP*s) may guide the recognition of various pathogens (70), initiating the TOLL-signaling pathway and leading to the production of antimicrobial peptides such as attacin (71), gloverin (72), lysozyme (73), and cecropin (74). Although we cannot establish causality between expression differences of immune genes and investment in reproduction or migration, immune gene evolution has been shown to be dynamic in migratory species in general (68) and may be of particular importance in V. cardui where several immune genes are uniquely present in multi-copy arrays (35).

In addition to immune response adjustments, efficient utilization of energy is of ultimate importance in migratory species. In insects, fat serves as the most efficient source for storage of energy (75,76) and lipids are indeed the main fuel for flight (77). Corroborating that, we observed that pathways associated with metabolism have critical roles in resource allocation trade-offs (71), we found that host plant availability variation resulted in differential expression of multiple genes associated with lipid and carbohydrate metabolism, for example, dihydrodiol dehydrogenase (*DHDH*) and dolihol kinase (*DOLK*) genes (79).

Our study highlights the central role of hormonal regulation, metabolism, and immunity in butterflies’ response to host plant availability. Although our data and approaches do not allow us to establish a causative association between host plant availability and investment in reproduction or migration per se, the gene expression analysis revealed a set of candidate genes that can be used to investigate the molecular underpinnings of the reproduction-migration tradeoff in more detail. These findings may extend beyond the classical oogenesis-flight syndrome, suggesting broader applicability in understanding synchronization between reproduction and migration. The next step will be to target key genes in the regulatory pathways detected here, with a particular focus on the ecdysone pathway. It should be noted that the trade-off between reproduction and migration in adult butterflies is likely not exclusively dictated by environmental cues encountered after emergence. As a complementary step, we therefore explored how differences in food plant availability and crowding affect the expression profiles across ontogenetic stages, from larvae to imagines.

### 4.2 Effects on gene expression by differences in food plant availability and rearing density during development

Both crowding and food resource availability have been shown to impact the timing of development and morphology of migratory insects, which in turn directly affect the flight response norms and migration propensity (80,81). Insect individuals experiencing starvation in general exhibit delayed development (33), prolonged larval stages and reduced body sizes (81), and these effects have also previously been observed in *V. cardui* (82). Similarly, negative associations between larval density and developmental rates have been established (28). This evidence arises from direct measurement of resulting phenotypic traits, while molecular mechanisms involved in the responses are less studied (but see e.g. 30). In our study, transcriptomic signatures of the response to periodic starvation and larval crowding confirmed differences in the activation of developmental genes and pathways in multiple organ systems. Differentially expressed genes were associated with ontology terms such as epithelial cell and dendrite development, sarcomere organization, angiogenesis, and hemocyte proliferation.

Gene expression profiles suggest that the food limitation treatment appeared to specifically affect neural development. Neural development and sensing are of particular importance for migratory insects, as they can largely influence the plasticity of the response to perceived environmental cues in adults (81). In our experiment, two candidate genes—odorant receptor 4 and esterase (odorant degrading protein)—were activated in adult individuals who had not experienced food stress as larvae, suggesting enhanced environmental sensing (84). Conversely, the downregulation of the gooseberry-neuro (*GsbN*) transcription factor suggests alterations in central nervous system development (85). We therefore consider these particular genes as key candidates for further investigation of how environmental cues are translated into behavioral responses.

The larval density experiment triggered differential expression of genes related development of reproductive systems, which is illustrated by ontology terms such as sperm individualization and gonad morphogenesis. Early changes in reproductive functions are particularly noteworthy in the context of the trade-off between reproduction and migration in the adult stage, a topic previously discussed in relation to the host plant experiment. These results underscore the role of the male reproductive system as well. Our result is in line with observations in other lepidopterans, such as that *Plodia interpunctella*, and *Mythimna separata*, which have been shown to have increased sperm production in response to crowding (86).

Alterations in metabolism at the molecular level appear to be a consistent response to all environmental cues tested in this study, as evidenced by the earlier-mentioned responses to host-plant presence in adult females, and the impacts of crowding and food scarcity across developmental stages and in adults. Notably, in the context of the starvation treatment, we identified over 200 differentially expressed genes associated with the ‘compound metabolic process’. On the individual gene level, we observed differential expression of the trehalose transporter (*Tret1*) in adults who experienced starvation during development. Trehalose is the primary sugar found in insect hemolymph, synthesized in the fat body and subsequently distributed by transporters (87). Alteration of expression of *Tret1* was also directly linked to active migration in insects (63). Among other candidate genes differentially expressed in the density experiment were the metabolic genes NADH dehydrogenase and several chitinases (86).

Since the analysis spanned multiple developmental stages, from larval instar III to recently emerged imagines, we gained insights into the temporal variation in gene expression and identified critical developmental periods where the effect was particularly pronounced. Notably, the most significant gene expression differences occurred during the larval stages in both experiments, especially in larval instar V—the final stage for *V. cardui* and a critical time point for responding to environmental cues before metamorphosis (89). Our findings highlight genes associated with programmed cell death, such as E3-type small ubiquitin-like modifier (*SUMO*) (90), and show enrichment in the JNK and Ras signaling pathways, essential for the tissue remodeling required during metamorphosis (91). In general, the last larval stage is accompanied by considerable physiological changes and environmental shifts during this period can therefore have particular importance for plastic responses (92).

## 5 CONCLUSIONS

In a host plant experiment designed to trigger the trade-off between migration and reproduction, we observed signatures of gene expression consistent with those expected for the oogenesis-flight syndrome and highlighted the crucial role of hormonal regulation. By subjecting larvae to different environmental cues, food abundance and larval crowding, we examined the early predisposition for migratory plasticity. This approach allowed us to closely track the timing of cue perception throughout development. Our findings revealed the peak of this response during the last larval stage, emphasizing the role of the genes involved in developmental regulation and metabolism. Furthermore, this led us to identify candidate genes and pathways that jointly contribute to the migratory syndrome.

## DATA AVAILABILITY AND BENEFIT-SHARING

Raw sequence data (RNA-seq) are available at the European Nucleotide Archive under XXXXXXXX. The scripts used to generate the analyses presented in the paper are archived on GitHub in the following repository: https://github.com/orgs/EBC-butterfly-genomics-team.

The benefit arising from the utilization of genetic resources in this study is provided through the sharing of the analysis scripts and code, which are freely available and accessible to the research community, consistent with applicable international and national regulations.

## AUTHOR CONTRIBUTIONS

NB contributed to the conception and design of the study. Lab work was performed by EP and LH with assistance from KN and AP. Formal analysis and visualization were performed by DS with assistance from VT. DS and NB wrote the manuscript. GT provided samples. NB supervised the study. Funding acquisition was handled by NB, GT, and RV. All authors reviewed and approved the final manuscript.

## ACKNOWLEDGEMENTS

Acknowledgments This work was funded by a research grant from the Swedish Research Council FORMAS (grant # 2019-00670 to N.B.) and The Swedish Collegium for Advanced Science (Natural Sciences Programme, Knut and Alice Wallenberg Foundation, Postdoc funding for D.S.). R.V. was supported by grant PID2022-139689NB-I00 (MICIU/ AEI/ 10.13039/501100011033 and ERDF, EU) and by grant 2021-SGR-00420 (Departament de Recerca i Universitats, Generalitat de Catalunya). G.T. was supported by grants PID2020-117739GA-I00 MCIN / AEI / 10.13039/ 501100011033, LINKA20399 from the CSIC iLink program and 2021-SGR-01334 (Departament de Recerca i Universitats, Generalitat de Catalunya). The authors acknowledge support from the National Genomics Infrastructure in Stockholm funded by Science for Life Laboratory, the Knut and Alice Wallenberg Foundation and the Swedish Research Council, and SNIC/Uppsala Multidisciplinary Center for Advanced Computational Science for assistance with massively parallel sequencing and access to the UPPMAX computational infrastructure. The computations were enabled by resources provided by the National Academic Infrastructure for Supercomputing in Sweden (NAISS) and the Swedish National Infrastructure for Computing (SNIC) in Uppsala, partially funded by the Swedish Research Council through grant agreements no. 2022-06725 and no. 2018-05973. We thank Jesper Boman and all members of the Backström lab for helpful discussions.

